# Ultra-High Multiplexing Enables Near-Full-Length 16S rRNA Gene Amplicon Sequencing of Over 1,200 Gut Microbiome Samples on a Single Nanopore Flow Cell

**DOI:** 10.64898/2026.08.29.747698

**Authors:** Charlie Hejlskov McPhillips, Eoghan Thomas Reilly, Gladys Stolberg-Mathieu, Klara Nielsen, Adam Duun Gottlieb, Gjorgji Madjarov, Henrik M. Roager, Dennis Sandris Nielsen, Lukasz Krych

## Abstract

Next-generation sequencing (NGS) of the prokaryotic 16S rRNA gene revolutionized gut microbiome research two decades ago. However, short read lengths remain an inherent limitation of platforms such as the widely used Illumina platforms (2 x 150-300 bp). Recent advances in Oxford Nanopore Technologies (ONT) flow cell chemistry (R10.4.1) have substantially improved sequencing accuracy. Combined with a custom multiple-primer strategy that comprehensively targets 16S rRNA gene variants to generate near-full-length amplicons, this approach enables read-by-read taxonomic classification, a feature not feasible with short-read sequencing platforms. Although our multiple-primer strategy could enable parallel sequencing of more than 18,000 samples (192 x 96), current flow cell capacity offers sufficient sequencing depth for approximately 1,000-1,500 samples. To validate the scalability and our per-read classification pipeline, we show that more than a thousand human fecal microbiome samples spiked with two bacterial strains (*Imtechella halotolerans* and *Allobacillus halotolerans*), not otherwise present in human fecal samples, can be successfully sequenced on a single flow cell, achieving a per-molecule error rate sufficient for direct per-read classification and at an adequate read depth for downstream analysis. This level of scalability significantly reduces per-sample costs, making the approach more accessible to a broader research community. To embrace these advancements, we have developed RubyRed, a pipeline that processes raw sequencing data and assigns taxonomic classifications on a per-read basis.

Using spike-in references (*I. halotolerans* and *A. halotolerans*), we demonstrate high mean single-read sequencing accuracy (99% and 98.9%, respectively), with the majority of reads exceeding the canonical threshold required for species-level taxonomic classification based on the 16S rRNA gene.

## Introduction

Taxonomic profiling is a foundational component in characterizing complex microbial communities like the gut microbiome. During the last two decades, amplicon sequencing of the 16S rRNA gene has been widely adopted for this purpose due to its scalability and cost-effectiveness. Early large-scale studies of the gut microbiome were driven by NGS short-read platforms, which made profiling accessible to a broad research community (Gehrig et al., 2022; Mahmoud et al., 2025).

Next-generation sequencing platforms have revolutionized microbial community studies by enabling highly parallelized amplicon sequencing at massive throughput. However, these technologies are constrained by inherently short read lengths, which typically range from 150 bp to approximately 550 bp, depending on the platform (e.g., up to 2 x 300 bp on Illumina MiSeq). Variation in achievable read length across platforms has been a major explanation for the limited standardization in studies utilizing 16S rRNA gene amplicon sequencing. Consequently, many protocols utilize specific hypervariable regions for amplification not only based on conserved primer-binding sites and the discriminatory power of variable regions, but also according to platform-specific read length limitations. For example, platforms capable of generating ~ 500 bp reads (2 x 300 bp) typically target the V3–V4 hypervariable regions, whereas platforms limited to 150 bp or 250 bp reads are often restricted to single-region protocols targeting V3 *or* V4 alone.

Advancements in full-length sequencing have revealed that intragenomic regions of the rRNA gene (e.g., 16S and 18S) have variations that are widespread and should not be ignored in microbiome profiling (Wang et al., 2023). By sequencing the full-length 16S rRNA gene, rather than short-read sequencing of variable regions, phylogenetic classification can be performed more accurately, even for low-frequency variants (Johnson et al., 2019). In addition, primer bias and difficulties in resolving closely related taxa can lead to incomplete or ambiguous community profiles (Wang et al., 2023; Agustinho et al., 2024). These limitations have motivated the development and adoption of sequencing strategies that provide both increased resolution and improved accuracy.

Long-read sequencing technologies, particularly those offered by Oxford Nanopore Technologies (ONT) and Pacific Biosciences (PacBio), have emerged as powerful tools for microbiome profiling. By enabling sequencing of near-full-length 16S rRNA genes and longer amplicons, these platforms substantially improve species-level classification and phylogenetic inference, resulting in more accurate representations of microbial community structure (Biada et al., 2025; Agustinho et al., 2024). PacBio has traditionally delivered highly accurate long reads, whereas early ONT-based amplicon sequencing was challenged by relatively high single-molecule error rates. However, ONT offers several major advantages, including substantially lower instrument costs, higher throughput, and reduced per-sample sequencing costs, making it an attractive alternative provided that single-molecule accuracy can be sufficiently improved.

Current ONT platforms report single-molecule accuracy on 16S amplicon sequencing of above 99% (Zhang et al., 2023). Furthermore, we have recently demonstrated that data generated using the R10.4.1 flow cell chemistry can be used to reconstruct complete, high-accuracy bacterial genomes and methylomes that are comparable to, and in some cases exceed, the accuracy achieved with PacBio sequencing (Soto-Serrano et al., 2024). Since the quality on the single molecule level has now exceeded the 16S rRNA gene similarity level generally necessary for species level classification (98.5%, Rodriguez et al., 2018), we have developed a novel pipeline, RubyRed, that performs read-by-read classification for rapid data analysis, making the need for actual operational taxonomic units (OTUs) obsolete.

In this study, we exploit recent advances in nanopore sequencing technology to demonstrate the scalability of ONT-based amplicon sequencing for profiling the human gut microbiome. Our previously reported protocol, which employs a multiple-primer strategy for comprehensive targeting of 16S rRNA gene variants combined with custom PCR barcoding, was integrated with ONT’s native barcoding kit. This approach theoretically enables pooling of up to 18,432 samples (192 x 96) within a single sequencing library. However, considering current throughput of PromethION flow cells, we show that an optimal balance between sequencing depth and throughput is achieved at approximately 1,200 samples per flow cell. By sequencing no more than 1,200 samples on a single PromethION flow cell (R10.4.1), we demonstrate that this strategy delivers sufficient depth and accuracy to comprehensively characterize complex microbial communities such as the human gut microbiome, highlighting its potential as a very cost-effective solution for large-scale taxonomic profiling.

## Results

### RubyRed processing pipeline filters and classifies

Sequencing of the library generated on average more than 6 million reads per barcode (NBC 1–16). The reads were processed using the RubyRed pipeline (Figure 3). Approximately 46% of reads were removed during primer and length filtering, as they either lacked the expected primer sequences or did not match the expected 16S rRNA gene length (V1–V9). An additional 3.6% of reads were discarded due to low average sequence quality (Phred score < 15).

To standardize sequencing depth across samples and reduce downstream computational costs, samples exceeding 30,000 reads per barcode were subsampled (user-adjustable optional step), removing approximately 16% of all reads. More than 2 million reads remained and were subsequently screened for chimeric sequences, resulting in a further 2.4% reduction. The remaining reads were then taxonomically classified, and the 5.1% that could not be assigned to a taxon were discarded. Assuming that reads removed during the subsampling step contain similar proportions of chimeric and unassigned sequences as the retained reads, a sequencing output of 6 million raw reads per barcode would yield approximately 2.5 million high-quality reads passing all processing steps and suitable for confident taxonomic assignment. The required sequencing depth ultimately depends on the experimental design and the expected complexity of the microbial community. As approximately 60% of raw reads are typically removed during the processing pipeline, it is worth noting that the real-time read count reported by MinKNOW can be used to estimate the expected number of high-quality, taxonomically assignable reads using the following equation:

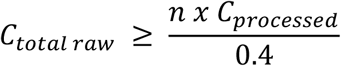

If a lower sequencing depth is sufficient for the intended analyses, sequencing can be terminated earlier. For example, when sequencing 96 samples in a full-plate format, a target depth of *C*_*processed*_ = 20,000 *reads* per sample for downstream analyses would require a total raw read count of *C*_*total raw*_ ≥ 4,800,000 reads. This approach enables users to stop the sequencing run once the required amount of data has been collected. This approach provides a cost-efficient strategy for processing multiple samples through multiplexing, substantially reducing the per-sample cost (Figure 2A). Following sequencing, the flow cell can be washed and reused for subsequent runs (e.g., using the EXP-WSH004 Flow Cell Wash Kit; Oxford Nanopore Technologies), further improving overall cost efficiency.

### Sequencing accuracy exceeds the 16S rRNA gene species-level classification threshold

Using the spike-in controls, we assessed the average sequencing error rate of the assigned reads, by BLASTing assigned reads against the reference sequences (Supplementary: table S2). BLAST percent identity (pident) scores showed that reads classified as either of the two spike-in strains, *I. halotolerans* and *A. halotolerans*, had mean identities of 99.0% and 98.9%, respectively; both exceeding the species-level classification threshold for 16S rRNA gene (Rodriguez et al., 2018) (Supplementary: table S3).

Across the sixteen barcodes, a total of 103,448 reads were assigned to *I. halotolerans* and 110,138 reads to *A. halotolerans*. Each species accounted for approximately 0.27% of the total classified reads across all barcodes. Of the reads assigned to *I. halotolerans* and *A. halotolerans*, 84.1% and 80.7%, respectively, had percent identity values above the classification threshold (Figure 4).

## Discussion

### Read depth defines scaling limits

High throughput 16S rRNA gene amplicon sequencing remains a cost effective and rapid tool for bacterial relative composition analysis across multiple samples. Short-read sequencing technologies, mainly Illumina, have been used extensively for the past two decades by many research groups. Although the near-full-length 16S rRNA reads allow for more detailed taxonomic classification, genus level analysis has often proved sufficient for basic conclusions on e.g., microbiome perturbations. Hence, short read sequencing was preferred due to its low cost per sample. The lower basecalling accuracy of early versions of ONT sequencing further discouraged the utilization of near-full-length reads of the 16S rRNA gene.

Recent advances in Oxford Nanopore Technologies (ONT) sequencing have substantially improved single-read accuracy. Combined with the increased throughput of PromethION flow cells, which can now generate up to 200 Gbp of sequence data per flow cell, this enables the simultaneous analysis of far more samples than is possible with MinION or GridION flow cells. By combining our previously described custom PCR barcoding strategy, first published in 2021 (Arildsen et al., 2021) and benchmarked against short-read sequencing technology (Hui et al., 2021), with ONT’s native barcoding kit, up to 18,432 individual samples can be analysed in parallel.

Naturally, with current PromethION’s flow cell (R 10.4.1), the throughput would result in a very shallow average read depth if processing the maximum number of samples, utilizing the full set of available barcodes (n = 18,432). In this study, sequencing of 1,209 samples achieved a minimum read depth of close to 10,000 reads per sample for the majority of samples (90%), after quality filtering through RubyRed (Figure 2B). Increasing the number of samples beyond 1,200 would likely reduce the average and minimum read depth, leading to a higher proportion of samples with low read depth. As flow cell capacity could be expected to increase in the future, the number of samples per flow cell is likely to increase accordingly, making this method future-proof.

Samples with reduced sequencing depth were most likely affected by errors during the pooling step. Such samples can be repooled and resequenced either separately or by adding them at a higher concentration to an existing library pool. When only a small number of samples are loaded onto a new or previously used flow cell, the required sequencing time is substantially shorter, and sufficient data may be generated within minutes to a few hours. This provides an additional advantage of real-time sequencing technology. In the present study, the entire Plate 5 library was repooled and resequenced on a new flow cell (data not shown).

### Per-read classification replaces clustering and denoising

Recent advances in ONT sequencing technology, particularly improvements in R10.4.1 flow cells and basecalling algorithms (Dorado v7.9.8), have dramatically reduced sequencing error rates. In parallel, increased computational power and optimized taxonomic classification algorithms now enable direct taxonomic classification of individual long reads without relying on OTU clustering or ASV inference (Johnson et al., 2019).

Historically, OTU clustering was introduced primarily to reduce the computational burden of taxonomic classification when sequencing throughput exceeded available computing resources. By clustering highly similar sequences prior to classification, millions of reads could be represented by a much smaller number of operational taxonomic units, substantially accelerating downstream analyses, albeit at the expense of sequence-level resolution (Nguyen et al., 2016).

The subsequent introduction of amplicon sequence variants (ASVs) aimed to preserve single-nucleotide differences by modelling and correcting sequencing errors (Callahan et al., 2017). While this represented a major advance for short-read sequencing, ASV inference remains dependent on statistical denoising of reads, where genuine biological variants and polymerase-derived errors may differ by few nucleotides. Consequently, the distinction between true low-abundance variants and amplification artefacts cannot always be resolved with certainty (Nearing et al., 2018).

In contrast, the high single-read accuracy of ONT sequencing enables direct read-by-read taxonomic classification of full-length 16S rRNA gene sequences. Rather than attempting to reconstruct putative biological sequence variants, each read is independently assigned to its most likely taxonomic origin based on the information contained within the entire gene. This approach avoids clustering or denoising while preserving all observed reads for downstream analyses, providing a simple, computationally efficient, and taxonomically focused framework that is well suited for routine microbiome profiling.

Nevertheless, RubyRed also supports conventional OTU clustering and ASV inference as optional downstream analyses for users who prefer these established approaches or require them for compatibility with existing workflows. OTU- and ASV-based feature tables can be generated for analyses that depend on sequence clustering, such as phylogeny-based diversity metrics (e.g. weighted and unweighted UniFrac) or direct comparison with previously published studies.

## Materials and Methods

### DNA library preparation and sequencing

Informed consent was obtained from all participants in the MOTILITY Mother-Child cohort (Stolberg-Mathieu et al., 2025). A total of 1,181 fecal samples (60 samples from mothers and 1,121 samples from infants) collected in the MOTILITY Mother-Child cohort (Stolberg-Mathieu et al., 2025) and mock communities (n = 28) were used for DNA extraction. Briefly, fecal material was homogenized 1:1 with Milli-Q water, aliquoted, and stored at −70°C until further analysis. Homogenates were thawed on ice and vortexed briefly. A minimum of 300-350 mg homogenate was transferred to a clean Eppendorf tube and spun down at 12000 rpm for 5 minutes. The supernatant was removed, and 125 mg of the remaining pellet and 20 µL ZymoBIOMICS Spike-in Control I (containing two alien bacterial strains; *Imtechella halotolerans* and *Allobacillus halotolerans*; Supplementary: Table S2) were transferred to a PowerBead Pro tube. Subsequent DNA extraction was carried out according to the QIAGEN DNA-extraction protocol but using a bead-beater (MP-FastPrep-24, MP Biomedicals) for 60 s at 4.5-6.5 m/s. DNA concentration was measured using Qubit dsDNA HS Assay Kits and diluted in Milli-Q water to 5 ng/µL per sample in 96-well plates. A two-step PCR approach (Hui et al., 2021) was then employed to amplify the 16S rRNA gene and barcode the amplicons. The first PCR was performed by mixing 5 µL of template DNA (5 ng/µL) with 12 µL of 2X PCRBIO Ultra Mix and 2 µL of primer mix (5 µM UMI_338ab_F/UMI_1391_R and UMI_27ab_F/UMI_1540_R; Supplementary: table S1) in a total reaction volume of 25 µL. PCR amplification was carried out on a thermocycler (SureCycler 8800, Agilent) using the following conditions: 95 °C for 5 min; 2 cycles of 95 °C for 20 s, 48 °C for 30 s, 65 °C for 10 s, and 72 °C for 45 s; followed by a final extension at 72 °C for 4 min. PCR products were subsequently purified using AMPure XP beads (Beckman Coulter, CA, USA) on an automated liquid-handling system according to the manufacturer’s instructions. Purified PCR products were then subjected to a second round of PCR to amplify near-full-length 16S rRNA gene amplicons and incorporate custom barcodes. For this step, 11 µL of purified PCR product was transferred directly from the magnetic plate into a new 96-well plate and mixed with 12 µL of 2X PCRBIO Ultra Mix and 2 µL of ONT UMI16S SET1 barcodes. Amplification was performed on a thermocycler (SureCycler 8800, Agilent) under the following conditions: 95 °C for 2 min; 33 cycles of 95 °C for 20 s, 55 °C for 20 s, and 72 °C for 40 s; followed by a final extension at 72 °C for 4 min. The resulting PCR products were assessed by agarose gel electrophoresis by mixing 8 µL of PCR product with 2 µL of 6X loading dye and running it in a 1.5% agarose gel in 0.5X TBE buffer for 45 min at 120 V. DNA bands were visualized following incubation with 0.8% ethidium bromide solution. Concentrations of amplified PCR products were measured using the Qubit 1X dsDNA HS Assay Kit on a Qubit fluorometer (Thermo Fisher Scientific, USA).

The PCR plates containing the amplified 16S gene-sequences were pooled into individual Eppendorf tubes at equimolar ratios, ensuring approximately 3 ng of each sample in the final pool. Pooled samples were purified using AMPure XP beads (Beckman Coulter Genomics, CA, USA) and subjected to native barcoding using the SQK-NBD114.24 protocol. Native-barcoded pools were subsequently combined into a single Eppendorf tube containing all samples (n > 1,200) and loaded onto a PromethION flow cell (Oxford Nanopore Technologies, R10.4.1) (Figure 1).

**Figure 1.**
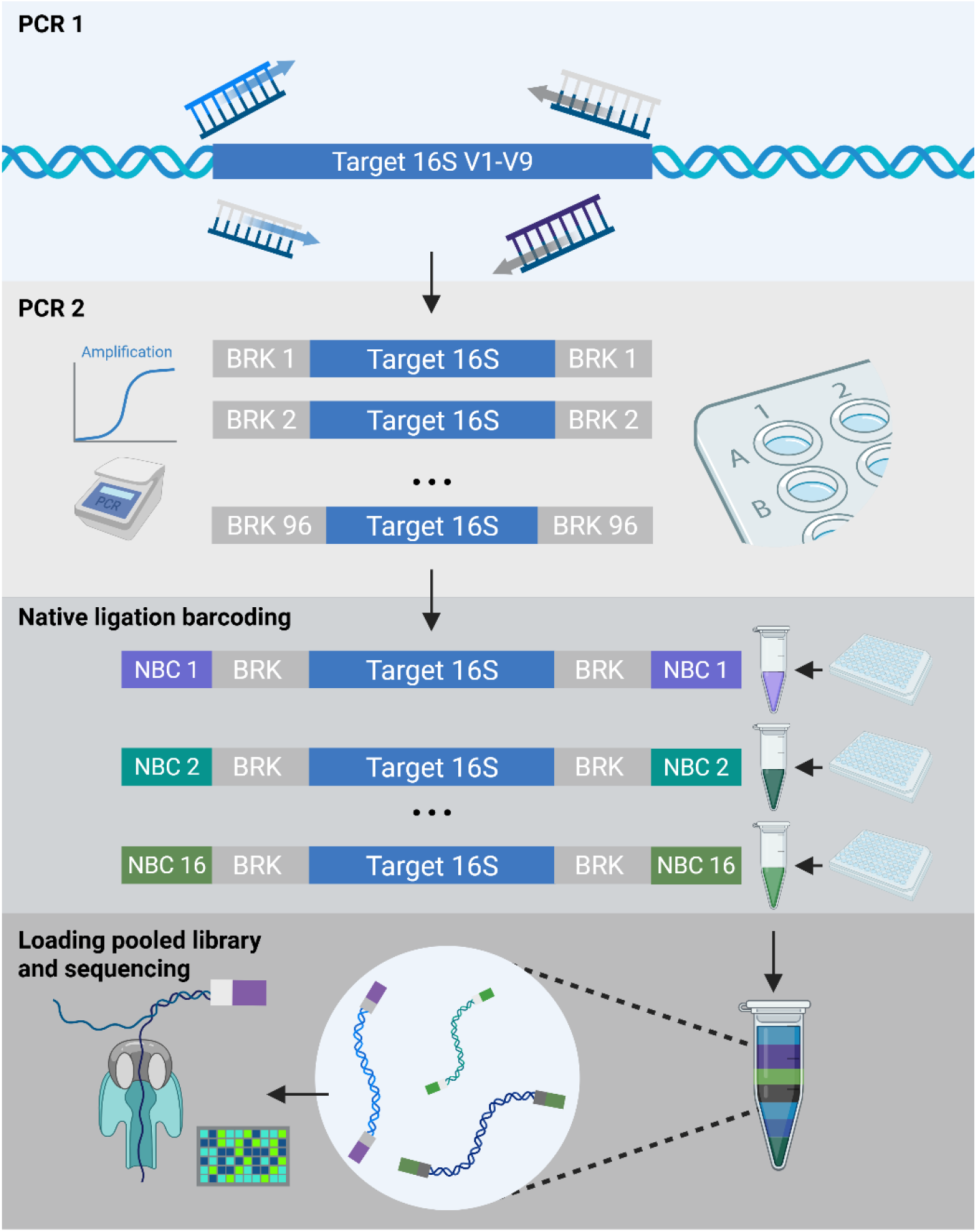
Method overview: Target 16S rRNA gene spanning the variable regions (V1-V9), are amplified in two cycles using four primers (27Fa/b, 338Fa/b, 1391R,1540R, S1). Near-full length 16S genes are then amplified and tagged with a barcode enabling downstream demultiplexing by PCR, in theory this could be done with the full-set of 192 barcodes. Each plate is pooled into a single Eppendorf, where native ligation barcoding adds a second identifier, ONT’s native barcode (NBC). After cleaning of the pooled samples, all the samples can be added to a single flow cell. Basecalling is performed by Dorado (v. 7.9.8) integrated into MinKNOW (v. 25.04.14); taxonomic classification was carried out through our custom read-by-read classifier RubyRed.

**Figure 2.**
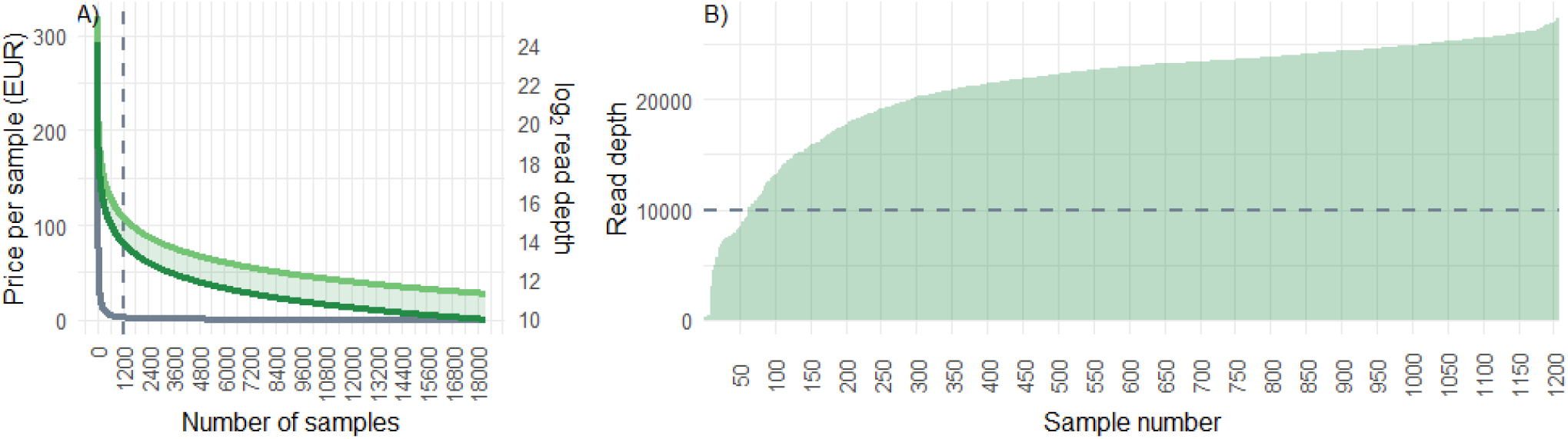
Read depth massively reduces operating costs, while still providing adequate sequencing depth for taxonomic classification. A) cost of running a single sample is around 3200 EUR. Custom multi-primer barcoding reduces this to a fraction of this by supporting the sequencing of over a thousand individual samples in a single flow cell (R 10.4.1). Green lines: log2 transformed read depth (light green: raw; dark green processed), grey line: price per sample. Dashed vertical line is the 1200 samples used to validate the scalability B) Read depth distribution of ordered samples; lowest read-count was found to be 9,771 reads. Rubyred has a default cut-off threshold at 30,000 reads per sample to lower the processing time.

**Figure 3.**
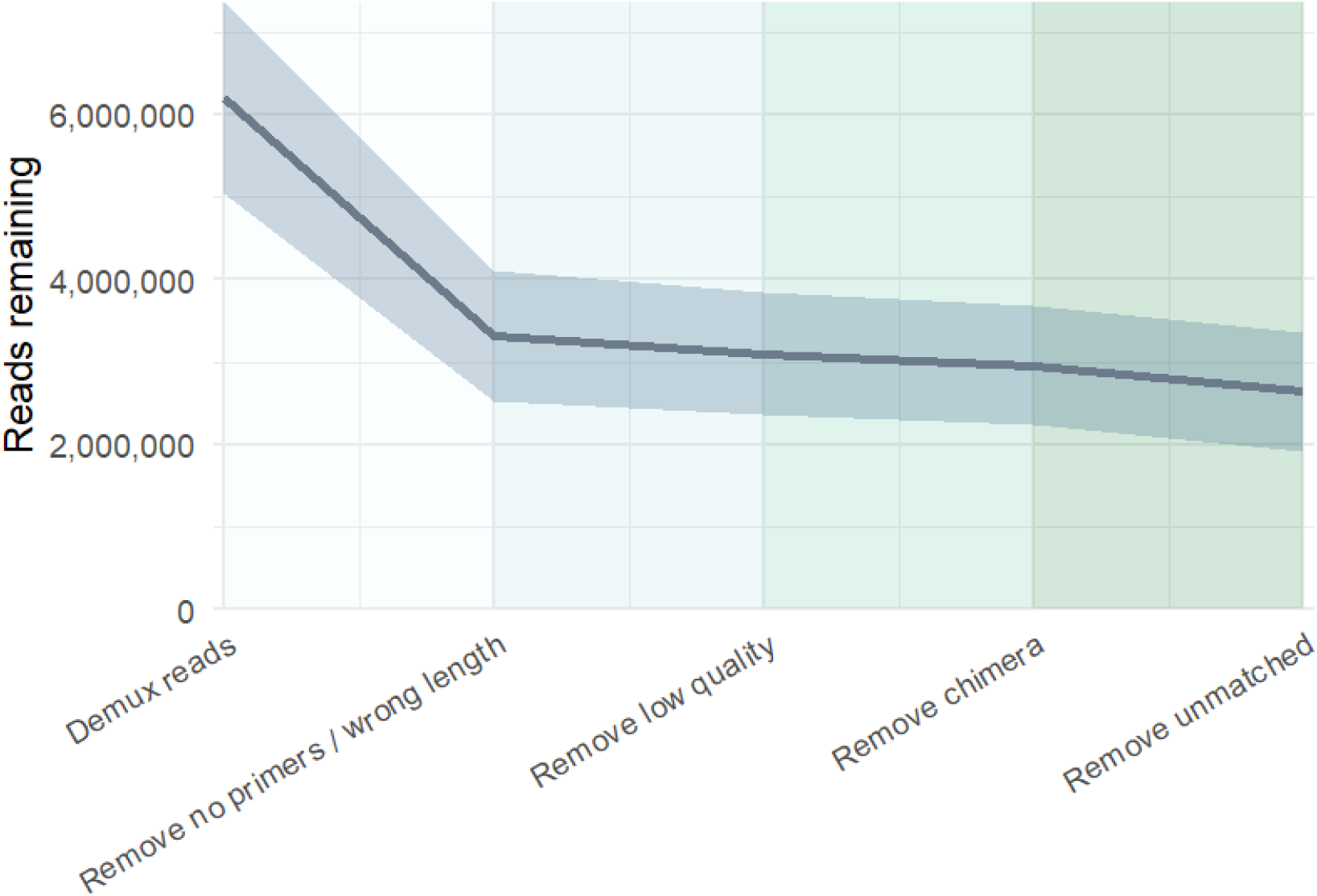
RubyRed pipeline processing steps (mean ±SD across barcodes). The pipeline sequentially filters raw reads by removing sequences lacking primers or with incorrect length, discarding low-quality reads, eliminating chimeric sequences, and matching reads to a reference database for taxonomic assignment. An average of 43% of reads across the 16 barcodes pass all processing steps and are retained for downstream analysis.

**Figure 4.**
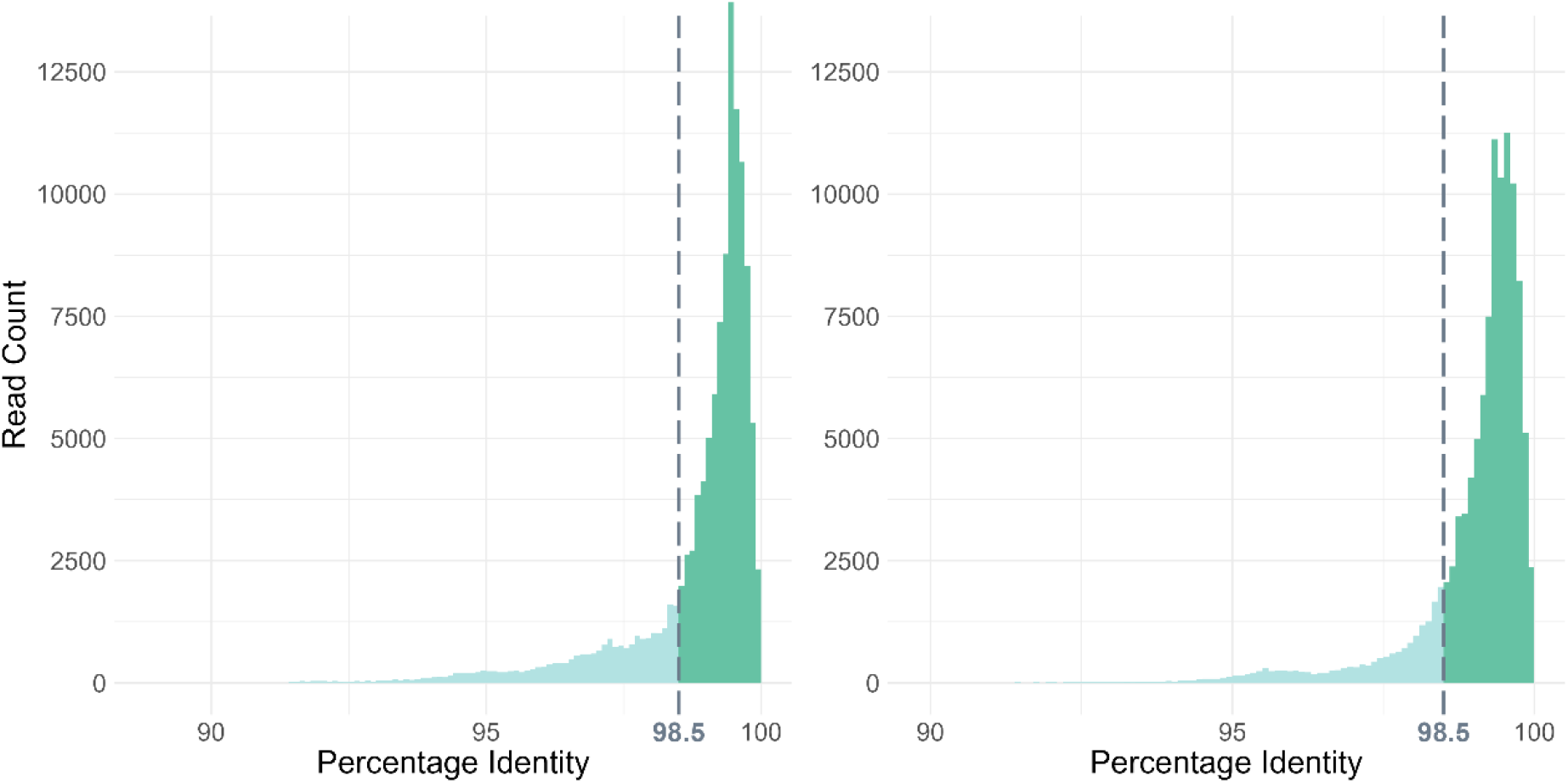
Distribution of reads assigned to the two spike-in strains, Allobacillus halotolerans (left) and Imtechella halotolerans (right). Mean sequencing accuracy is 98.9% and 99%, respectively. The dashed line indicates the canonical threshold for species-level classification based on the 16S gene.

## Data Analysis

### Basecalling and first level demultiplexing

Basecalling on the raw pod5 files was performed in real-time using (Dorado v7.9.8) in super-accurate mode. First level demultiplexing was performed using Dorado (v7.9.8) integrated into MinKNOW (v25.04.14). Demultiplexed barcodes (separation by PCR plates) were subjected for further analysis using a custom developed pipeline, RubyRed.

### Data processing and read-by-read classification

A custom bioinformatics pipeline, RubyRed (https://github.com/reillyeo/RubyRed), was developed to perform high-throughput, read-by-read taxonomic classification directly from Oxford Nanopore FASTQ files. The pipeline includes a custom demultiplexing tool, Torchlex (Schack et al., 2024), which performs barcode assignment with higher speed and accuracy than the Guppy demultiplexer (Oxford Nanopore Technologies) in our benchmark datasets (data not shown). Following demultiplexing, reads undergo primer trimming with cutadapt (Martin, 2011), quality filtering with chopper (De Coster & Rademakers, 2023), subsampling with seqkit (Shen et al., 2024), and FASTA conversion with vsearch (Rognes et al., 2016), before merging reads and importing them as a single sequence artifact into QIIME2 (Bolyen et al., 2019). After utilizing QIIME2’s inbuilt vsearch and rescript plugins for chimera removal and strand reorientation, respectively, RubyRed assigns taxonomy using one of three classification methods: sklearn (default), consensus-vsearch, or consensus-blast, with optional abundance filtering to reduce noise. The default database used for reference-based chimera filtering and strand reorientation, as well as both reference-based classification (vsearch, BLAST) and training of sklearn’s machine learning-based classifier, is the MIMt-16s rRNA database (release version M2c_26_03) (Cabezas et al., 2024), a database of amplicon sequences compiled from NCBI RefSeq and curated to maximize quality and eliminate redundancy. However, several other 16S rRNA gene databases are available, as well as databases for other rRNA amplicons (18S, ITS, 28S). The pipeline outputs taxonomically annotated feature tables and representative sequences for downstream analysis. Other default parameters include a minimum average read quality of 15, acceptable length after primer removal between 800 and 1600 bp, and a subsampling depth of 30,000 reads, all of which are user-customizable.

## Conclusion

Advances in ONT flow cell chemistry have enabled per-read classification, which combined with a multiple-primer strategy, has immense potential for analyzing a large number of samples in parallel. To accommodate this, we have developed a pipeline that processes raw reads and classifies them on a per-read basis. Although our custom barcoding system enables demultiplexing of more than 18,000 samples, current sequencing capacity is better suited to fewer samples to provide sufficient processed read depth for downstream analysis. To validate the pipeline and demonstrate its scalability, this study provides proof of concept by sequencing more than 1,200 samples on a single flow cell. As flow cell chemistry and sequencing capacity are expected to improve in the future, we demonstrate a future-proof approach in analyzing vast numbers of gut microbiomes effectively and simultaneously.

## Acknowledgment

We would like to thank the experts from G+D Netcetera, Zypressenstrasse 71, 8004 Zurich, Switzerland for support help in development of Torchlex. We thank the children and families for participating in the MOTILITY Mother-Child cohort.

## Data availability

The primary outcome of the MOTILITY study has not been published yet. Hence raw data is not available at present. Of note, the aim of the present manuscript is to show proof-of-concept of the possibilities for massive parallel sequencing using the developed protocol. However, we will happily provide access to positive and negative controls, as well as *Imtechella halotolerans* and *Allobacillus halotolerans* counts by emailing the corresponding author Lukasz Krych.

