## Supplemental Table S1 for "Ultra-High Multiplexing Enables Near-Full-Length 16S rRNA Gene Amplicon Sequencing of Over 1,200 Gut Microbiome Samples on a Single Nanopore Flow Cell"

Table S3: BLAST pident on reads assigned to either of the spike-in strains

| Barcode ID | *Allobacillus halotolerans* | | *Imtechella halotolerans* | |
| --- | --- | --- | --- | --- |
|  | Identity, % | Number of reads | Identity, % | Number of reads |
| barcode01 | 98.8 | 3933 | 99.0 | 5201 |
| barcode02 | 99.0 | 5585 | 99.1 | 5087 |
| barcode03 | 98.9 | 4276 | 99.0 | 4451 |
| barcode04 | 98.9 | 7772 | 99.1 | 8882 |
| barcode05 | 98.9 | 9032 | 98.9 | 8623 |
| barcode06 | 99.0 | 9319 | 99.1 | 8902 |
| barcode07 | 98.9 | 7757 | 99.1 | 5993 |
| barcode08 | 98.9 | 4598 | 99.0 | 4225 |
| barcode09 | 98.9 | 6718 | 99.1 | 6677 |
| barcode10 | 98.9 | 9371 | 99.0 | 8705 |
| barcode11 | 98.9 | 3145 | 99.0 | 2707 |
| barcode12 | 98.8 | 4669 | 99.0 | 4157 |
| barcode13 | 98.8 | 3117 | 99.0 | 2720 |
| barcode14 | 98.9 | 4063 | 99.0 | 3072 |
| barcode15 | 99.0 | 9529 | 99.1 | 8729 |
| barcode16 | 99.1 | 17254 | 99.1 | 15317 |
| Mean | 98.9 | 6883 | 99.0 | 6465 |
