## Supplemental Table S2 for "Ultra-High Multiplexing Enables Near-Full-Length 16S rRNA Gene Amplicon Sequencing of Over 1,200 Gut Microbiome Samples on a Single Nanopore Flow Cell"

Table S2: 16S gene sequences from Allobacillus halotolerans and Imtechella halotolerans used to assess sequencing error

>Allobacillus_halotolerans

GAGAGTTTGATCCTGGCTCAGGATGAACGCTGGCGGCGTGCCTAATACATGCAAGTCGAGCGCGGGAAGCAGACAGCGTCCCTTCGGGGACAATGTCTGTGGAACGAGCGGCGGACGGGTGAGTAACACGTGGGCAACCTGCCTGCAAGACTGGGATAACCCCGGGAAACCGGGGCTAATACCGGATGATGACATGAATCGCATGATTCATGTTTGAAAGATGGCCTTTGTGCTATCACTTGCAGATGGGCCCGCGGCGCATTAGCTAGTTGGTGGGGTAAGGGCCTACCAAGGCAACGATGCGTAGCCGACCTGAGAGGGTGATCGGCCACACTGGGACTGAGACACGGCCCAGACTCCTACGGGAGGCAGCAGTAGGGAATCTTCCGCAATGGACGAAAGTCTGACGGAGCAACGCCGCGTGAGTGAAGAAGGTCCTCGGATCGTAAAGCTCTGTTGTTAGGGAAGAACAAGTACCGTTCGAATAGGGCGGTACCTTGACGGTACCTAACCAGAAAGCCACGGCTAACTACGTGCCAGCAGCCGCGGTAATACGTAGGTGGCAAGCGTTATCCGGAATTATTGGGCGTAAAGCGCGCGCAGGCGGTTTCTTAAGTCTGATGTGAAAGCCCACGGCTTAACCGTGGAGGGTCATTGGAAACTGGGGAACTTGAGTACAGAAGAGGAGAGTGGAATTCCACGTGTAGCGGTGAAATGCGTAGAGATGTGGAGGAACACCAGTGGCGAAGGCGACTCTCTGGTCTGTAACTGACGCTGAGGCGCGAAAGCGTGGGTAGCAAACAGGATTAGATACCCTGGTAGTCCACGCCGTAAACGATGAGTGCTAGGTGTTAGGGGTTCCAACCCTTAGTGCTGCAGTTAACGCAATAAGCACTCCGCCTGGGGAGTACGGCCGCAAGGCTGAAACTCAAAGGAATTGACGGGGGCCCGCACAAGCGGTGGAGCATGTGGTTTAATTCGACGCAACGCGAAGAACCTTACCAGGTCTTGACATCCCGCTGACCGGCCTAGAGATAGGCCTTCCCTTCGGGGCAGCGGTGACAGGTGGTGCATGGTTGTCGTCAGCTCGTGTCGTGAGATGTTGGGTTAAGTCCCGTAACGAGCGCAACCCTTGATCTTAGTTGCCAGCATTTAGTTGGGCACTCTAAGGTGACTGCCGGTGACAAACCGGAGGAAGGTGGGGATGACGTCAAATCATCATGCCCCTTATGACCTGGGCTACACACGTGCTACAATGGATGGTACAGAGGGCAGCGAGACCGTGAGGTTTAGCCAATCCCTTAAAGCCATTCTCAGTTCGGATTGCAGGCTGCAACTCGCCTGTATGAAGCCGGAATCGCTAGTAATCGCGGATCAGCATGCCGCGGTGAATACGTTCCCGGGCCTTGTACACACCGCCCGTCACACCACGAGAGTTGGCAACACCCGAAGTCGGTGGGGTAACCTTTATGGAGCCAGCCGCCGAAGGTGGGGCCAATGATTGGGGTGAAGTCGTAACAAGGTAGCCGTATCGGAAGGTGCGGCTGGATCACCTCCTTTCTA

>Allobacillus_halotolerans

GAGAGTTTGATCCTGGCTCAGGATGAACGCTGGCGGCGTGCCTAATACATGCAAGTCGAGCGCGGGAAGCAGACAGCGTCCCTTCGGGGACAATGTCTGTGGAACGAGCGGCGGACGGGTGAGTAACACGTGGGCAACCTACCTGCAAGACTGGGATAACCCCGGGAAACCGGGGCTAATACCGGATGATGACGTGAATCGCATGATTCATGTTTGAAAGATGGCCTTTGTGCTATCACTTGCAGATGGGCCCGCGGCGCATTAGCTAGTTGGTGGGGTAAGAGCCTACCAAGGCAACGATGCGTAGCCGACCTGAGAGGGTGATCGGCCACACTGGGACTGAGACACGGCCCAGACTCCTACGGGAGGCAGCAGTAGGGAATCTTCCGCAATGGACGAAAGTCTGACGGAGCAACGCCGCGTGAGTGAAGAAGGTCCTCGGATCGTAAAGCTCTGTTGTTAGGGAAGAACAAGTACCGTTCGAATAGGGCGGTACCTTGACGGTACCTAACCAGAAAGCCACGGCTAACTACGTGCCAGCAGCCGCGGTAATACGTAGGTGGCAAGCGTTATCCGGAATTATTGGGCGTAAAGCGCGCGCAGGCGGTTTCTTAAGTCTGATGTGAAAGCCCACGGCTTAACCGTGGAGGGTCATTGGAAACTGGGGAACTTGAGTACAGAAGAGGAGAGTGGAATTCCACGTGTAGCGGTGAAATGCGTAGAGATGTGGAGGAACACCAGTGGCGAAGGCGACTCTCTGGTCTGTAACTGACGCTGAGGCGCGAAAGCGTGGGTAGCAAACAGGATTAGATACCCTGGTAGTCCACGCCGTAAACGATGAGTGCTAGGTGTTAGGGGTTCCAACCCTTAGTGCTGCAGTTAACGCAATAAGCACTCCGCCTGGGGAGTACGGCCGCAAGGCTGAAACTCAAAGGAATTGACGGGGGCCCGCACAAGCGGTGGAGCATGTGGTTTAATTCGACGCAACGCGAAGAACCTTACCAGGTCTTGACATCCCGCTGACCGGCCTAGAGATAGGCCTTCCCTTCGGGGCAGCGGTGACAGGTGGTGCATGGTTGTCGTCAGCTCGTGTCGTGAGATGTTGGGTTAAGTCCCGTAACGAGCGCAACCCTTGATCTTAGTTGCCAGCATTAAGTTGGGCACTCTAAGGTGACTGCCGGTGACAAACCGGAGGAAGGTGGGGATGACGTCAAATCATCATGCCCCTTATGACCTGGGCTACACACGTGCTACAATGGATGGTACAGAGGGCAGCGAGACCGTGAGGTTTAGCCAATCCCTTAAAGCCATTCTCAGTTCGGATTGCAGGCTGCAACTCGCCTGTATGAAGCCGGAATCGCTAGTAATCGCGGATCAGCATGCCGCGGTGAATACGTTCCCGGGCCTTGTACACACCGCCCGTCACACCACGAGAGTTGGCAACACCCGAAGTCGGTGGGGTAACCTTTATGGAGCCAGCCGCCGAAGGTGGGGCCAATGATTGGGGTGAAGTCGTAACAAGGTAGCCGTATCGGAAGGTGCGGCTGGATCACCTCCTTTCTA

>Allobacillus_halotolerans

GAGAGTTTGATCCTGGCTCAGGATGAACGCTGGCGGCGTGCCTAATACATGCAAGTCGAGCGCGGGAAGCAGACAGCGTCCCTTCGGGGACAATGTCTGTGGAACGAGCGGCGGACGGGTGAGTAACACGTGGGCAACCTACCTGCAAGACTGGGATAACCCCGGGAAACCGGGGCTAATACCGGATGATGACGTGAATCGCATGATTCATGTTTGAAAGATGGCCTTTGTGCTATCACTTGCAGATGGGCCCGCGGCGCATTAGCTAGTTGGTGGGGTAAGAGCCTACCAAGGCAACGATGCGTAGCCGACCTGAGAGGGTGATCGGCCACACTGGGACTGAGACACGGCCCAGACTCCTACGGGAGGCAGCAGTAGGGAATCTTCCGCAATGGACGAAAGTCTGACGGAGCAACGCCGCGTGAGTGAAGAAGGTCCTCGGATCGTAAAGCTCTGTTGTTAGGGAAGAACAAGTACCGTTCGAATAGGGCGGTACCTTGACGGTACCTAACCAGAAAGCCACGGCTAACTACGTGCCAGCAGCCGCGGTAATACGTAGGTGGCAAGCGTTATCCGGAATTATTGGGCGTAAAGCGCGCGCAGGCGGTTTCTTAAGTCTGATGTGAAAGCCCACGGCTTAACCGTGGAGGGTCATTGGAAACTGGGGAACTTGAGTACAGAAGAGGAGAGTGGAATTCCACGTGTAGCGGTGAAATGCGTAGAGATGTGGAGGAACACCAGTGGCGAAGGCGACTCTCTGGTCTGTAACTGACGCTGAGGCGCGAAAGCGTGGGTAGCAAACAGGATTAGATACCCTGGTAGTCCACGCCGTAAACGATGAGTGCTAGGTGTTAGGGGTTCCAACCCTTAGTGCTGCAGTTAACGCAATAAGCACTCCGCCTGGGGAGTACGGCCGCAAGGCTGAAACTCAAAGGAATTGACGGGGGCCCGCACAAGCGGTGGAGCATGTGGTTTAATTCGACGCAACGCGAAGAACCTTACCAGGTCTTGACATCCCGCTGACCGGCCTAGAGATAGGCCTTCCCTTCGGGGCAGCGGTGACAGGTGGTGCATGGTTGTCGTCAGCTCGTGTCGTGAGATGTTGGGTTAAGTCCCGTAACGAGCGCAACCCTTGATCTTAGTTGCCAGCATTAAGTTGGGCACTCTAAGGTGACTGCCGGTGACAAACCGGAGGAAGGTGGGGATGACGTCAAATCATCATGCCCCTTATGACCTGGGCTACACACGTGCTACAATGGATGGTACAGAGGGCAGCGAGACCGTGAGGTTTAGCCAATCCCTTAAAGCCATTCTCAGTTCGGATTGCAGGCTGCAACTCGCCTGTATGAAGCCGGAATCGCTAGTAATCGCGGATCAGCATGCCGCGGTGAATACGTTCCCGGGCCTTGTACACACCGCCCGTCACACCACGAGAGTTGGCAACACCCGAAGTCGGTGGGGTAACCTTTATGGAGCCAGCCGCCGAAGGTGGGGCCAATGATTGGGGTGAAGTCGTAACAAGGTAGCCGTATCGGAAGGTGCGGCTGGATCACCTCCTTTCTA

>Allobacillus_halotolerans

GAGAGTTTGATCCTGGCTCAGGATGAACGCTGGCGGCGTGCCTAATACATGCAAGTCGAGCGCGGGAAGCAGACAGCGTCCCTTCGGGGACAATGTCTGTGGAACGAGCGGCGGACGGGTGAGTAACACGTGGGCAACCTACCTGCAAGACTGGGATAACCCCGGGAAACCGGGGCTAATACCGGATGATGACGTGAATCGCATGATTCATGTTTGAAAGATGGCCTTTGTGCTATCACTTGCAGATGGGCCCGCGGCGCATTAGCTAGTTGGTGGGGTAAGAGCCTACCAAGGCAACGATGCGTAGCCGACCTGAGAGGGTGATCGGCCACACTGGGACTGAGACACGGCCCAGACTCCTACGGGAGGCAGCAGTAGGGAATCTTCCGCAATGGACGAAAGTCTGACGGAGCAACGCCGCGTGAGTGAAGAAGGTCCTCGGATCGTAAAGCTCTGTTGTTAGGGAAGAACAAGTACCGTTCGAACAGGGCGGTACCTTGACGGTACCTAACCAGAAAGCCACGGCTAACTACGTGCCAGCAGCCGCGGTAATACGTAGGTGGCAAGCGTTATCCGGAATTATTGGGCGTAAAGCGCGCGCAGGCGGTTTCTTAAGTCTGATGTGAAAGCCCACGGCTTAACCGTGGAGGGTCATTGGAAACTGGGGAACTTGAGTACAGAAGAGGAGAGTGGAATTCCACGTGTAGCGGTGAAATGCGTAGAGATGTGGAGGAACACCAGTGGCGAAGGCGACTCTCTGGTCTGTAACTGACGCTGAGGCGCGAAAGCGTGGGTAGCAAACAGGATTAGATACCCTGGTAGTCCACGCCGTAAACGATGAGTGCTAGGTGTTAGGGGTTCCAACCCTTAGTGCTGCAGTTAACGCAATAAGCACTCCGCCTGGGGAGTACGGCCGCAAGGCTGAAACTCAAAGGAATTGACGGGGGCCCGCACAAGCGGTGGAGCATGTGGTTTAATTCGACGCAACGCGAAGAACCTTACCAGGTCTTGACATCCCGCTGACCGGCCTAGAGATAGGCCTTACCTTCGGGGCAGCGGTGACAGGTGGTGCATGGTTGTCGTCAGCTCGTGTCGTGAGATGTTGGGTTAAGTCCCGTAACGAGCGCAACCCTTGATCTTAGTTGCCAGCATTAAGTTGGGCACTCTAAGGTGACTGCCGGTGACAAACCGGAGGAAGGTGGGGATGACGTCAAATCATCATGCCCCTTATGACCTGGGCTACACACGTGCTACAATGGATGGTACAGAGGGCAGCGAGACCGTGAGGTTTAGCCAATCCCTTAAAGCCATTCTCAGTTCGGATTGCAGGCTGCAACTCGCCTGTATGAAGCCGGAATCGCTAGTAATCGCGGATCAGCATGCCGCGGTGAATACGTTCCCGGGCCTTGTACACACCGCCCGTCACACCACGAGAGTTGGCAACACCCGAAGTCGGTGGGGTAACACAATACGTGAGCCAGCCGCCGAAGGTGGGGCCAATGATTGGGGTGAAGTCGTAACAAGGTAGCCGTATCGGAAGGTGCGGCTGGATCACCTCCTTTCTA

>Allobacillus_halotolerans

GAGAGTTTGATCCTGGCTCAGGATGAACGCTGGCGGCGTGCCTAATACATGCAAGTCGAGCGCGGGAAGCAGACAGCGTCCCTTCGGGGACAATGTCTGTGGAACGAGCGGCGGACGGGTGAGTAACACGTGGGCAACCTACCTGCAAGACTGGGATAACCCCGGGAAACCGGGGCTAATACCGGATGATGACGTGAATCGCATGATTCATGTTTGAAAGATGGCCTTTGTGCTATCACTTGCAGATGGGCCCGCGGCGCATTAGCTAGTTGGTGGGGTAAGAGCCTACCAAGGCAACGATGCGTAGCCGACCTGAGAGGGTGATCGGCCACACTGGGACTGAGACACGGCCCAGACTCCTACGGGAGGCAGCAGTAGGGAATCTTCCGCAATGGACGAAAGTCTGACGGAGCAACGCCGCGTGAGTGAAGAAGGTCCTCGGATCGTAAAGCTCTGTTGTTAGGGAAGAACAAGTACCGTTCGAATAGGGCGGTACCTTGACGGTACCTAACCAGAAAGCCACGGCTAACTACGTGCCAGCAGCCGCGGTAATACGTAGGTGGCAAGCGTTATCCGGAATTATTGGGCGTAAAGCGCGCGCAGGCGGTTTCTTAAGTCTGATGTGAAAGCCCACGGCTTAACCGTGGAGGGTCATTGGAAACTGGGGAACTTGAGTACAGAAGAGGAGAGTGGAATTCCACGTGTAGCGGTGAAATGCGTAGAGATGTGGAGGAACACCAGTGGCGAAGGCGACTCTCTGGTCTGTAACTGACGCTGAGGCGCGAAAGCGTGGGTAGCAAACAGGATTAGATACCCTGGTAGTCCACGCCGTAAACGATGAGTGCTAGGTGTTAGGGGTTCCAACCCTTAGTGCTGCAGTTAACGCAATAAGCACTCCGCCTGGGGAGTACGGCCGCAAGGCTGAAACTCAAAGGAATTGACGGGGGCCCGCACAAGCGGTGGAGCATGTGGTTTAATTCGACGCAACGCGAAGAACCTTACCAGGTCTTGACATCCCGCTGACCGGCCTAGAGATAGGCCTTCCCTTCGGGGCAGCGGTGACAGGTGGTGCATGGTTGTCGTCAGCTCGTGTCGTGAGATGTTGGGTTAAGTCCCGTAACGAGCGCAACCCTTGATCTTAGTTGCCAGCATTAAGTTGGGCACTCTAAGGTGACTGCCGGTGACAAACCGGAGGAAGGTGGGGATGACGTCAAATCATCATGCCCCTTATGACCTGGGCTACACACGTGCTACAATGGATGGTACAGAGGGCAGCGAGACCGTGAGGTTTAGCCAATCCCTTAAAGCCATTCTCAGTTCGGATTGCAGGCTGCAACTCGCCTGTATGAAGCCGGAATCGCTAGTAATCGCGGATCAGCATGCCGCGGTGAATACGTTCCCGGGCCTTGTACACACCGCCCGTCACACCACGAGAGTTGGCAACACCCGAAGTCGGTGGGGTAACCTTTATGGAGCCAGCCGCCGAAGGTGGGGCCAATGATTGGGGTGAAGTCGTAACAAGGTAGCCGTATCGGAAGGTGCGGCTGGATCACCTCCTTTCTA

>Allobacillus_halotolerans

GAGAGTTTGATCCTGGCTCAGGATGAACGCTGGCGGCGTGCCTAATACATGCAAGTCGAGCGCGGGAAGCAGACAGCGTCCCTTCGGGGACAATGTCTGTGGAACGAGCGGCGGACGGGTGAGTAACACGTGGGCAACCTACCTGCAAGACTGGGATAACCCCGGGAAACCGGGGCTAATACCGGATGATGACGTGAATCGCATGATTCATGTTTGAAAGATGGCCTTTGTGCTATCACTTGCAGATGGGCCCGCGGCGCATTAGCTAGTTGGTGGGGTAAGAGCCTACCAAGGCAACGATGCGTAGCCGACCTGAGAGGGTGATCGGCCACACTGGGACTGAGACACGGCCCAGACTCCTACGGGAGGCAGCAGTAGGGAATCTTCCGCAATGGACGAAAGTCTGACGGAGCAACGCCGCGTGAGTGAAGAAGGTCCTCGGATCGTAAAGCTCTGTTGTTAGGGAAGAACAAGTGCCGTTCGAACAGGGCGGTACCTTGACGGTACCTAACCAGAAAGCCACGGCTAACTACGTGCCAGCAGCCGCGGTAATACGTAGGTGGCAAGCGTTATCCGGAATTATTGGGCGTAAAGCGCGCGCAGGCGGTTTCTTAAGTCTGATGTGAAAGCCCACGGCTTAACCGTGGAGGGTCATTGGAAACTGGGGAACTTGAGTACAGAAGAGGAGAGTGGAATTCCACGTGTAGCGGTGAAATGCGTAGAGATGTGGAGGAACACCAGTGGCGAAGGCGACTCTCTGGTCTGTAACTGACGCTGAGGCGCGAAAGCGTGGGTAGCAAACAGGATTAGATACCCTGGTAGTCCACGCCGTAAACGATGAGTGCTAGGTGTTAGGGGTTCCAACCCTTAGTGCTGCAGTTAACGCAATAAGCACTCCGCCTGGGGAGTACGGCCGCAAGGCTGAAACTCAAAGGAATTGACGGGGGCCCGCACAAGCGGTGGAGCATGTGGTTTAATTCGACGCAACGCGAAGAACCTTACCAGGTCTTGACATCCCGCTGACCGGCCTAGAGATAGGCCTTCCCTTCGGGGCAGCGGTGACAGGTGGTGCATGGTTGTCGTCAGCTCGTGTCGTGAGATGTTGGGTTAAGTCCCGTAACGAGCGCAACCCTTGATCTTAGTTGCCAGCATTAAGTTGGGCACTCTAAGGTGACTGCCGGTGACAAACCGGAGGAAGGTGGGGATGACGTCAAATCATCATGCCCCTTATGACCTGGGCTACACACGTGCTACAATGGATGGTACAGAGGGCAGCGAGACCGTGAGGTTTAGCCAATCCCTTAAAGCCATTCTCAGTTCGGATTGCAGGCTGCAACTCGCCTGTATGAAGCCGGAATCGCTAGTAATCGCGGATCAGCATGCCGCGGTGAATACGTTCCCGGGCCTTGTACACACCGCCCGTCACACCACGAGAGTTGGCAACACCCGAAGTCGGTGGGGTAACCTTTATGGAGCCAGCCGCCGAAGGTGGGGCCAATGATTGGGGTGAAGTCGTAACAAGGTAGCCGTATCGGAAGGTGCGGCTGGATCACCTCCTTTCTA

>Imtechella_halotolerans

AAGAGTTTGATCCTGGCTCAGGATGAACGCTAGCGGCAGGCCTAACACATGCAAGTCGAGGGGTAGAGGTTACTTGTAACCTTGAGACCGGCGCACGGGTGCGTAACGCGTATGCAATCTACCTTGTACAGAGGGATAGCCCAGAGAAATTTGGATTAATACCTCATAGTGTATAGATATGGCCTCATATTTATACTAAAGCTTCGGCGGTACAAGATGAGCATGCGTCCCATTAGCTAGTTGGTATGGTAACGGCATACCAAGGCAACGATGGGTAGGGGTCCTGAGAGGGAGATCCCCCACACTGGTACTGAGACACGGACCAGACTCCTACGGGAGGCAGCAGTGAGGAATATTGGGCAATGGTCGGAAGACTGACCCAGCCATGCCGCGTGCAGGAAGACGGCCCTATGGGTTGTAAACTGCTTTTGTATGGGAAGAATAAGGCCTACGTGTAGGTTGATGACGGTACCATGCGAATAAGCATCGGCTAACTCCGTGCCAGCAGCCGCGGTAATACGGAGGATGCAAGCGTTATCCGGAATCATTGGGTTTAAAGGGTCCGTAGGCGGGCTTGTAAGTCAGAGGTGAAAGTTTGCGGCTCAACCGTAAAATTGCCTTTGATACTGCTAGTCTTGAATAATTGTGAAGTGGTTAGAATATGTAGTGTAGCGGTGAAATGCATAGATATTACATAGAATACCGATTGCGAAGGCAGATCACTAACAATTTATTGACGCTGATGGACGAAAGCGTGGGGAGCGAACAGGATTAGATACCCTGGTAGTCCACGCCGTAAACGATGGATACTAGCTGTTCGGCGCAAGCTGAGTGGCTAAGCGAAAGTGATAAGTATCCCACCTGGGGAGTACGTTCGCAAGAATGAAACTCAAAGGAATTGACGGGGGCCCGCACAAGCGGTGGAGCATGTGGTTTAATTCGATGATACGCGAGGAACCTTACCAGGGCTTAAATGTAGATTGACAGGTTTAGAGATAGACTTTCCTTCGGGCAATTTACAAGGTGCTGCATGGTTGTCGTCAGCTCGTGCCGTGAGGTGTCAGGTTAAGTCCTATAACGAGCGCAACCCCTGTGGTTAGTTACCAGCACATAATGGTGGGGACTCTAGCCAGACTGCCGGTGCAAACCGTGAGGAAGGTGGGGATGACGTCAAATCATCACGGCCCTTACGTCCTGGGCCACACACGTGCTACAATGGCAAGTACAGAGAGCAGCCACTGGGTGACCAGGAGCGAATCTATAAAGCTTGTCTCAGTTCGGATCGAAGTCTGCAACTCGACTTCGTGAAGCTGGAATCGCTAGTAATCGGATATCAGCCATGATCCGGTGAATACGTTCCCGGGCCTTGTACACACCGCCCGTCAAGCCATGGAAGCTGGGGGTACCTGAAGTCGGTGACCGCAAGGAGCTGCCTAGGGTAAAACTAGTAACTGGGGCTAAGTCGTAACAAGGTAGCCGTACCGGAAGGTGCGGCTGGAACACCTCCTTTCTA

>Imtechella_halotolerans

AAGAGTTTGATCCTGGCTCAGGATGAACGCTAGCGGCAGGCCTAACACATGCAAGTCGAGGGGTAGAGGTTACTTGTAACCTTGAGACCGGCGCACGGGTGCGTAACGCGTATGCAATCTACCTTGTACAGAGGGATAGCCCAGAGAAATTTGGATTAATACCTCATAGTGTATAGATATGGCCTCATATTTATACTAAAGCTTCGGCGGTACAAGATGAGCATGCGTCCCATTAGCTAGTTGGTATGGTAACGGCATACCAAGGCAACGATGGGTAGGGGTCCTGAGAGGGAGATCCCCCACACTGGTACTGAGACACGGACCAGACTCCTACGGGAGGCAGCAGTGAGGAATATTGGGCAATGGTCGGAAGACTGACCCAGCCATGCCGCGTGCAGGAAGACGGCCCTATGGGTTGTAAACTGCTTTTGTATGGGAAGAATAAGGCCTACGTGTAGGTTGATGACGGTACCATGCGAATAAGCATCGGCTAACTCCGTGCCAGCAGCCGCGGTAATACGGAGGATGCAAGCGTTATCCGGAATCATTGGGTTTAAAGGGTCCGTAGGCGGGCTTGTAAGTCAGAGGTGAAAGTTTGCGGCTCAACCGTAAAATTGCCTTTGATACTGCTAGTCTTGAATAATTGTGAAGTGGTTAGAATATGTAGTGTAGCGGTGAAATGCATAGATATTACATAGAATACCGATTGCGAAGGCAGATCACTAACAATTTATTGACGCTGATGGACGAAAGCGTGGGGAGCGAACAGGATTAGATACCCTGGTAGTCCACGCCGTAAACGATGGATACTAGCTGTTCGGCGCAAGCTGAGTGGCTAAGCGAAAGTGATAAGTATCCCACCTGGGGAGTACGTTCGCAAGAATGAAACTCAAAGGAATTGACGGGGGCCCGCACAAGCGGTGGAGCATGTGGTTTAATTCGATGATACGCGAGGAACCTTACCAGGGCTTAAATGTAGATTGACAGGTTTAGAGATAGACTTTCCTTCGGGCAATTTACAAGGTGCTGCATGGTTGTCGTCAGCTCGTGCCGTGAGGTGTCAGGTTAAGTCCTATAACGAGCGCAACCCCTGTGGTTAGTTACCAGCACATAATGGTGGGGACTCTAGCCAGACTGCCGGTGCAAACCGTGAGGAAGGTGGGGATGACGTCAAATCATCACGGCCCTTACGTCCTGGGCCACACACGTGCTACAATGGCAAGTACAGAGAGCAGCCACTGGGTGACCAGGAGCGAATCTATAAAGCTTGTCTCAGTTCGGATCGAAGTCTGCAACTCGACTTCGTGAAGCTGGAATCGCTAGTAATCGGATATCAGCCATGATCCGGTGAATACGTTCCCGGGCCTTGTACACACCGCCCGTCAAGCCATGGAAGCTGGGGGTACCTGAAGTCGGTGACCGCAAGGAGCTGCCTAGGGTAAAACTAGTAACTGGGGCTAAGTCGTAACAAGGTAGCCGTACCGGAAGGTGCGGCTGGAACACCTCCTTTCTA

>Imtechella_halotolerans

AAGAGTTTGATCCTGGCTCAGGATGAACGCTAGCGGCAGGCCTAACACATGCAAGTCGAGGGGTAGAGGTTACTTGTAACCTTGAGACCGGCGCACGGGTGCGTAACGCGTATGCAATCTACCTTGTACAGAGGGATAGCCCAGAGAAATTTGGATTAATACCTCATAGTGTATAGATATGGCCTCATATTTATACTAAAGCTTCGGCGGTACAAGATGAGCATGCGTCCCATTAGCTAGTTGGTATGGTAACGGCATACCAAGGCAACGATGGGTAGGGGTCCTGAGAGGGAGATCCCCCACACTGGTACTGAGACACGGACCAGACTCCTACGGGAGGCAGCAGTGAGGAATATTGGGCAATGGTCGGAAGACTGACCCAGCCATGCCGCGTGCAGGAAGACGGCCCTATGGGTTGTAAACTGCTTTTGTATGGGAAGAATAAGGCCTACGTGTAGGTTGATGACGGTACCATGCGAATAAGCATCGGCTAACTCCGTGCCAGCAGCCGCGGTAATACGGAGGATGCAAGCGTTATCCGGAATCATTGGGTTTAAAGGGTCCGTAGGCGGGCTTGTAAGTCAGAGGTGAAAGTTTGCGGCTCAACCGTAAAATTGCCTTTGATACTGCTAGTCTTGAATAATTGTGAAGTGGTTAGAATATGTAGTGTAGCGGTGAAATGCATAGATATTACATAGAATACCGATTGCGAAGGCAGATCACTAACAATTTATTGACGCTGATGGACGAAAGCGTGGGGAGCGAACAGGATTAGATACCCTGGTAGTCCACGCCGTAAACGATGGATACTAGCTGTTCGGCGCAAGCTGAGTGGCTAAGCGAAAGTGATAAGTATCCCACCTGGGGAGTACGTTCGCAAGAATGAAACTCAAAGGAATTGACGGGGGCCCGCACAAGCGGTGGAGCATGTGGTTTAATTCGATGATACGCGAGGAACCTTACCAGGGCTTAAATGTAGATTGACAGGTTTAGAGATAGACTTTCCTTCGGGCAATTTACAAGGTGCTGCATGGTTGTCGTCAGCTCGTGCCGTGAGGTGTCAGGTTAAGTCCTATAACGAGCGCAACCCCTGTGGTTAGTTACCAGCACATAATGGTGGGGACTCTAGCCAGACTGCCGGTGCAAACCGTGAGGAAGGTGGGGATGACGTCAAATCATCACGGCCCTTACGTCCTGGGCCACACACGTGCTACAATGGCAAGTACAGAGAGCAGCCACTGGGTGACCAGGAGCGAATCTATAAAGCTTGTCTCAGTTCGGATCGAAGTCTGCAACTCGACTTCGTGAAGCTGGAATCGCTAGTAATCGGATATCAGCCATGATCCGGTGAATACGTTCCCGGGCCTTGTACACACCGCCCGTCAAGCCATGGAAGCTGGGGGTACCTGAAGTCGGTGACCGCAAGGAGCTGCCTAGGGTAAAACTAGTAACTGGGGCTAAGTCGTAACAAGGTAGCCGTACCGGAAGGTGCGGCTGGAACACCTCCTTTCTA
