## Supplemental Table S3 for "Ultra-High Multiplexing Enables Near-Full-Length 16S rRNA Gene Amplicon Sequencing of Over 1,200 Gut Microbiome Samples on a Single Nanopore Flow Cell"

Table S1: Near-full-length 16S rRNA gene was amplified using a primer mix containing: 27Fa/b, 338Fa/b, 1391 R and 1540R. (Y. Hui et al., 2021)

| 27Fa/b | 5’- GTCTCGTGGG CTCGGNNNNN NNNNNNNNNN AGAGTTTGATYMTGGCTYAG −3’  5’- GTCTCGTGGG CTCGGNNNNN NNNNNNNNNN AGGGTTCGATTCTGGCTCAG −3’ |
| --- | --- |
| 338Fa/b | 5’- GTCTCGTGGG CTCGGNNNNN NNNNNNNNNN ACWCCTACGGGWGGCAGCAG −3’  5’- GTCTCGTGGG CTCGGNNNNN NNNNNNNNNN GACTCCTACGGGAGGCWGCAG −3’ |
| 1391 R | 5’- GTCTCGTGGG CTCGGNNNNN NNNNNNNNNN GACGGGCGGTGTGTRCA −3’ |
| 1540 R | 5’- GTCTCGTGGG CTCGGNNNNN NNNNNNNNNN TACGGYTACCTTGTTACGACT -3’ |
